# Molecular bases for the loss of type VI secretion system activity during enteroaggregative *E. coli* experimental evolution

**DOI:** 10.1101/2025.02.08.637259

**Authors:** Boris Taillefer, Jonas B. Desjardins, Eric Cascales

## Abstract

The type VI secretion system (T6SS) is a nanoweapon deployed by Gram-negative to inject effectors into target cells and hence involved in pathogenesis and bacterial competition. While T6SS gene clusters are found in all recently isolated commensals or pathogenic *Escherichia coli* strains, they are absent from classical laboratory strains. These *E. coli* strains, which have been used since decades for bacterial genetics, were usually grown in pure cultures, suggesting that T6SS might have been lost during evolution in absence of competitors. Here, we conducted a 640-generation experimental evolution by passaging the enteroaggregative *E. coli* (EAEC) 17-2 strain under controlled competition conditions against susceptible or immune recipient cells. EAEC T6SS activity was almost abolished when grown in the presence of immune recipients, while no difference with the ancestral strain was observed for EAEC grown in the presence of susceptible cells. Whole genome sequencing of 18 clones identified adaptative mutations responsible for the loss of T6SS activity, including mutations of the T6SS promoter and within genes encoding T6SS subunits, and revealed a novel regulatory mechanism involving RfaH-dependent antitermination. We further found that the RfaH binding site *ops* element is present in T6SS gene clusters of several species, suggesting a conserved RfaH-dependent regulation of T6SSs across enterobacteria. This work exemplifies the power of experimental evolution to understand T6SS genetic adaptation and to unravel new players for its function.

## INTRODUCTION

The type VI secretion system (T6SS) is a nanoweapon that delivers effectors into prokaryotic and eukaryotic cells, participating to interbacterial competition and virulence (1). The T6SS belongs to the broad family of contractile injection systems, which use a spring-like mechanism to propel an inner tube loaded with effectors. The T6SS is comprised of 14 core proteins that assemble two subcomplexes with distinct evolutionarily histories. In Proteobacteria, the membrane complex is constituted of the TssJLM proteins that recruits and anchors a cytoplasmic phage-like tubular structure. This tubular structure is constituted of a baseplate composed of TssEFGK wedges polymerized around the VgrG-PAAR spike complex, which serves as assembly platform for the inner tube wrapped by the contractile tube. Once assembled, the T6SS sheath contracts, propelling an effector-loaded needle towards the target cell. The sheath proteins are then disassembled and recycled by the dedicated ClpV ATPase (2).

Effectors delivered by the T6SS are either released in the environment, such as metallophores that chelate metals, or directly injected into target cells. We distinguish toxins with eukaryotic-specific (e.g., actin cross-linking), antibacterial-specific (e.g., peptidoglycan hydrolases, FtsZ-, 23S RNA- or EF-Tu-targeting ADP-ribosyltransferases), and trans-kingdom activities (e.g., DNases, proteases, phospholipases, NADH glycosylhydrolases) (3). T6SS^+^ bacteria species delivering antibacterial toxins protect themselves by the production of immunity proteins that usually bind and inhibit their cognate toxins.

T6SS genes are clustered in 20-70-kb large operons that are comprised of genes encoding the structural subunits of the apparatus, the toxins with the cognate immunities, and accessory proteins (4–8). These accessory proteins might be transcriptional or post-translational regulators, chaperones, adaptor proteins or proteins that facilitate the assembly or the mechanism of action of the T6SS. T6SS gene clusters are widespread in Gram-negative bacteria with an overrepresentation in Bacteroidetes and Proteobacteria (6,9–11). It is not only found in pathogens, but also in environmental species or commensal and symbionts strains. T6SS gene clusters are also abundant in pathogenic or commensal *Escherichia coli* species including all pathovar types or clinical isolates and categorize in the T6SS*^i^*-1, -2 and -3 families (12). Strikingly they are absent of all classical laboratory strains of *E. coli* such as MC4100, MG1655, W3110 or DH5α. Most of these strains derive from strains W1485 and 58-161 from Joshua Lederberg and Edward Tatum, which themselves originate from strain NCTC86 or K-12 isolated by Theodor Escherich in 1885, or J.E. Blair in 1922, respectively (13). Interestingly, the genome sequence of strain NCTC86 showed that it carries one copy of a T6SS gene cluster, typical of the T6SS*^i^*-2 group found in *E. coli* strains (14). The observation that *E. coli* laboratory strains are devoid of T6SS genes whereas original strains from which they derive and recently isolated strains carry one or several copies of clusters raised the question of how T6SS genes were lost upon years or decades of pure subcultures in rich media (15).

To reveal the mechanisms underlying loss of T6SS, we conducted an experimental evolution of the enteroaggregative *E. coli* (EAEC) strain 17-2 in a medium in which the Sci1 T6SS is active. This approach, which consists of subjecting a population to serial passages in a specific and controlled condition, allows to identify mutations that are responsible for a fitness gain or for adaptation (16). Here, we passaged the EAEC strain for 640 generations in the presence of susceptible or immune cells. Whereas the T6SS activity of the population evolved with susceptible cells was comparable to the ancestral strain, the T6SS activity of the population evolved with immune cells was significantly decreased, with a loss of 99% of the activity compared to the ancestral strain. To identify the genetic changes responsible for decrease or lost activity, we sequenced 18 clones with intermediate and null T6SS phenotypes identifying single-nucleotide polymorphisms (SNPs) in metabolic pathways (*glpK*, *rne*, *rpoC*, and *aqpZ*), in Sci1 T6SS promoter and genes (*tssM*, *tssK* and *tssF*), as well as in the *rfaH* and *csgE* genes. While the metabolic mutations correspond to adaptation to the medium, genetically reconstructed strains of T6SS and *rfaH* mutations confirmed their importance for T6SS activity. We then defined the molecular basis for the loss of activity of the T6SS and delineated the impact of the RfaH antiterminator in T6SS regulation.

## RESULTS

### Experimental design

To study the EAEC 17-2 T6SS Sci1 adaptation in the absence of target cell to eliminate, we first engineered a strain carrying an insertion of a kanamycin cassette in replacement of the *lacZ* gene (17-2 Δ*lacZ*Ω*kan*) for selection. This strain was then serially passaged for a total of approximatively 640 generations in sedimentation conditions in Sci1 T6SS inducing medium (SIM, M9 supplemented with glycerol and 10% LB) in the presence of susceptible W3110 *E. coli* K-12 cells or of immune EAEC 17-2 cells. Competition conditions were carried out for 10 generations, followed by a selection of the 17-2 Δ*lacZ*Ω*kan* strain on selective (kanamycin) X-Gal agar plates and archived prior to being subjected to a new passage (**Figure 1**).

**Figure 1.**
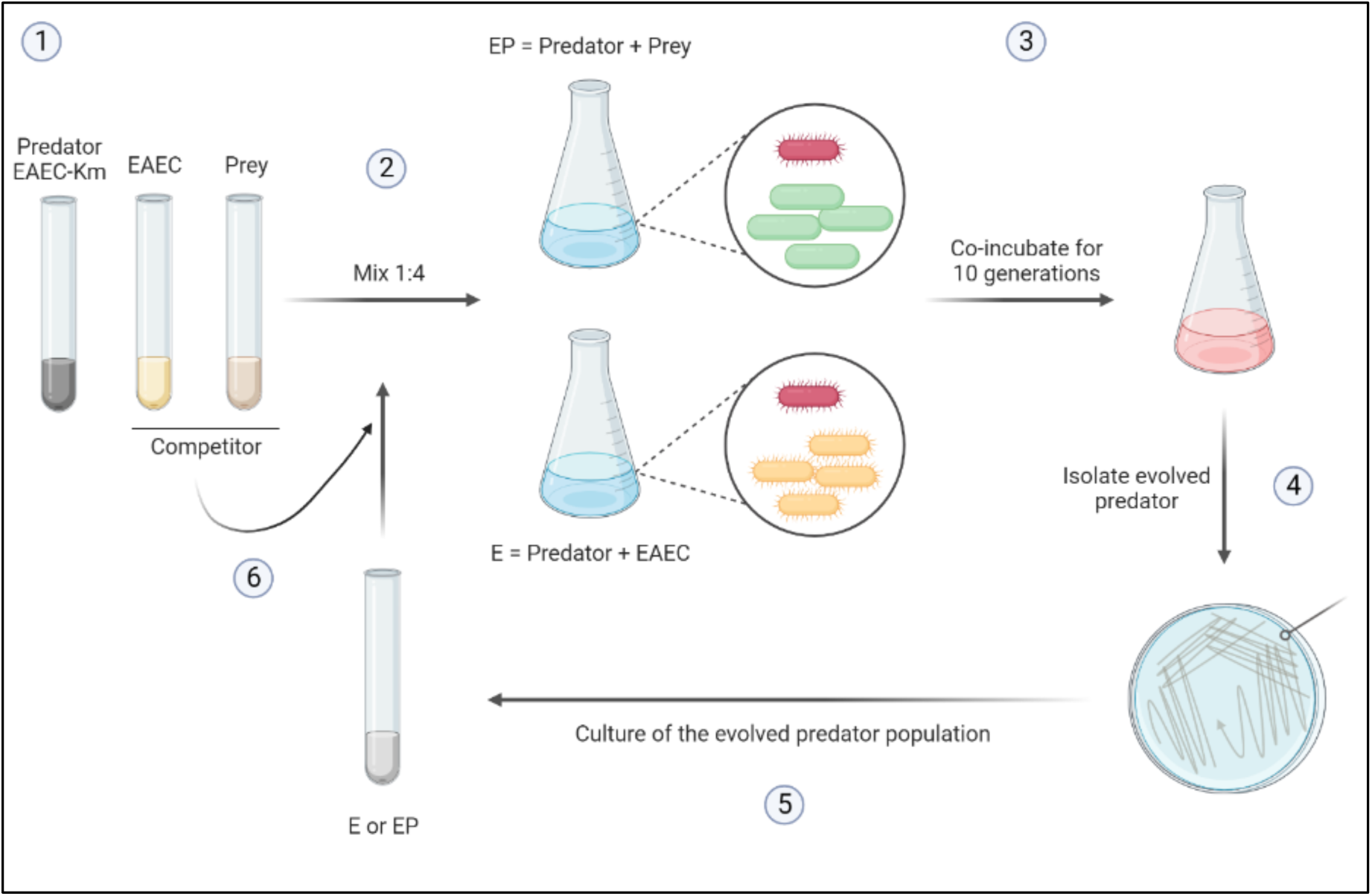
Schematic representation of the experimental evolution design. **(1)** The ancestral EAEC Δ*lacZ*Ω*kan* (EAEC-Km, T6SS^+^) strain and the susceptible (W3110, prey) and immune (EAEC) recipient were grown overnight from a single-colony. **(2)** EAEC-Km was mixed with W3110 (EP condition) or with EAEC (E condition) in a 1:4 ratio and **(3)** co-cultured in SIM at room temperature without agitation for 10 generations in sedimentation conditions. **(4)** The evolved EAEC-Km was isolated from the coculture by selection on kanamycin-LB plates, **(5)** resuspended in a liquid culture and **(6)** subjected to a new passage. The experiment was repeated 64 times (∼ 640 generations) to yield the EP640 and E640 populations. Created with Biorender.

### Experimental evolution in presence of immune cells led to a significant decrease of T6SS activity

The antibacterial activity of the resulting EAEC populations, named hereafter EP640 (evolved with W3110 recipient) and E640 (evolved with EAEC 17-2 recipient), were tested against the susceptible W3110 strain, and compared to the antibacterial activity of the ancestral strain or its Δ*sci1*derivative (deleted of the *sci1* T6SS gene cluster). **Figure 2A** shows that the EP640 population exhibited an antibacterial activity similar to that of the ancestral strain. In contrast, the E640 population was significantly impaired for T6SS activity, with a decrease of ∼ 99% compared to the ancestral strain (**Figure 2A**). These results indicate that most EAEC cells evolved with immune cells have lost T6SS activity, whereas this activity was conserved when evolved with susceptible cells, demonstrating that the T6SS provides a fitness advantage in the presence of sensible target cells.

**Figure 2.**
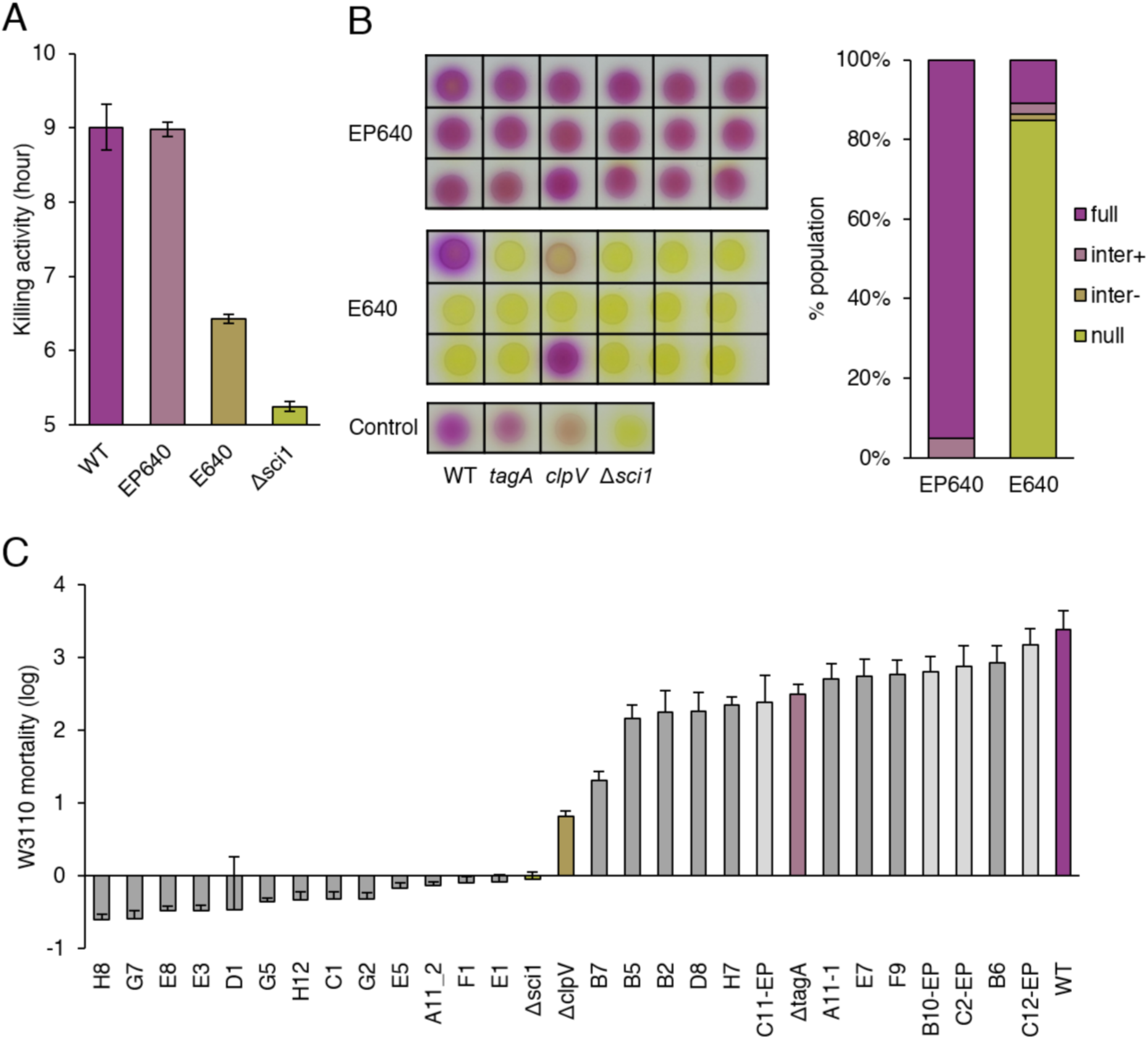
Loss of T6SS activity during experimental evolution of EAEC with immune cells. **A.** Competition assay between the EAEC ancestral (WT) strains or evolved E640 or EP640 populations or the T6SS-deletion mutant Δ*sci1* and the susceptible W3110 strain. W3110 mortality was measured by the SGK method. Data are the mean of three independent replicates. **B.** Representative antibacterial activity of selected clones from the EP640 (*n*=144) and E640 (*n*=192) populations using the LAGA assay (left). The ancestral EAEC (WT) and Δ*tagA*, Δ*clpV* and Δ*sci1* derivative strains were used as a control for full, intermediate plus, intermediate minus and null T6SS activity, respectively. The color intensity of CPRG (yellow to purple) correlates with the antibacterial activity. Clone phenotype percentage from EP640 and E640 population with wild-type, intermediate or null T6SS activity (left). This screening was performed on in a single replicate **C.** Antibacterial activity of selected evolved clones (light grey bars EP640, *n*=4; dark grey bars E640, *n*=22) using the SGK method. The ancestral strain (WT) and derivatives Δ*tagA*, Δ*clpV* and Δ*sci1*strains (colored bars) were used as controls. Data are the mean of at least three independent biological replicates.

To better characterize the EP640 and E640 populations, the antibacterial activity of 150-200 independent clones isolated from each population was assessed. Among the EP640 population, 97% of the 144 isolated clones presented a T6SS activity comparable to that of the ancestral strain, and 3% displayed a slightly reduced activity (**Figure 2B**). Conversely, ∼85% of the 192 E640 clones had no measurable T6SS activity, and 4% presented significantly reduced activity (**Figure 2B**). Twenty-six clones, including the 9 from E640 and 4 from EP640 population with intermediate T6SS activity, and 13 from E640 population with null activity, were selected for further analysis. Using a quantitative competition assay, we confirmed the intermediate antibacterial activity of the 9 E640 selected clones, with 4 presenting an activity between the WT and Δ*tagA* strains, and 5 others between the Δ*tagA* and Δc*lpV* mutant strains, and the absence of antibacterial activity for the 13 other E640 clones (**Figure 2C**). To test whether the absence of antibacterial activity is not linked to an impaired metabolism, the growth rate of all selected clones was measured. Although some clones displayed a reduced growth compared to the ancestral strain (longer lag time or lower biomass), only three of them presented significant growth defects and were not included for further analyses (**Figure S1**).

### Whole genome sequencing of evolved clones revealed mutations in genes involved in T6SS function

To understand how the phenotypes observed are linked to genetic changes, the ancestral strain and 18 evolved clones were subjected to whole genome sequencing using the Illumina technology. The genomes of selected clones were aligned and compared to the ancestral strain using the MAUVE software. The identified mutations, listed in **Table 1**, and illustrated in **Figure S2**, can be classified in three categories: metabolic adaptations (*glpK*, *rne*, *rpoC*, and *aqpZ* encoding a glycerol transporter, a ribonuclease, the RNA polymerase and the aquaporin Z, respectively); T6SS mutations (*sci1* promoter, *tssF*, *tssK* and *tssM* genes); other functions (*rfaH* and *csgE*). Mutations detected in evolved clones were either base substitutions (SNPs) or deletions (gap). We found 2 major lineages with metabolic adaptations [*rpoC*] and [*glpK*, *aqpZ*] (**Table S3).** The metabolic adaptation in *rne* gene was acquired independently in both lineages. These lineages appeared independently in the E640 and EP640 populations, in agreement with the fact that these metabolic mutations are adaptative mutations basically arising during experimental evolution in minimal media (17–20).

**Table 1.**
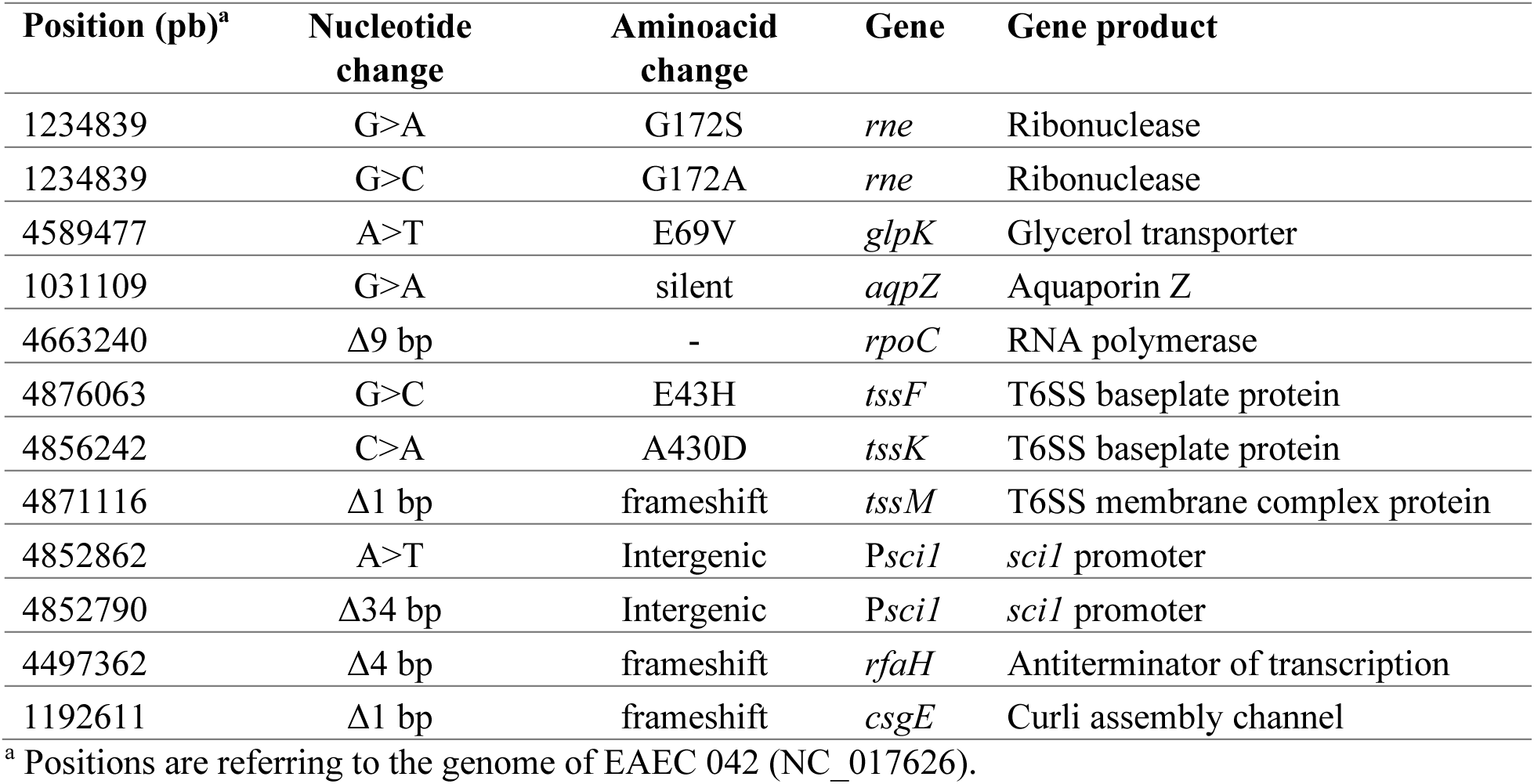
Mutations appeared during the EE in the selected evolved clones.

### Experimental evolution on minimal medium led to metabolic adaptation

As mentioned above, the metabolic adaptation mutations were already identified in independent studies where *E. coli* strains were evolved in starved conditions or in glycerol synthetic media (17). The D73K substitution in the glycerol-3-phosphate transporter GlpK, close to the E69 position identified here, increases GlpK catalytic activity and improve glycerol catabolism (17,19). The 9-bp deletion in the *rpoC* gene is identical to that found during evolution in glycerol (17) and is suggested to impact the global transcriptome in order to optimize growth in starved conditions (21). The synonymous mutation found in the *aqpZ* gene corresponds to residue A188 located in the loop E, which is involved in the substrate specificity (22–24). The next residue, R189, is important for water permeability (25). The substitutions changes the GCG codon for the less frequent GCA codon, and it is proposed that such a change in the bias codon may improve the overall fitness (26,27). Finally, two different mutations were found in the *rne* gene, producing a distinct substitution in the same amino acid (G172S ou G172A). The G172S substitution was already identified in *E. coli* MG1655 grown in minimal media supplemented with glycerol (17) and has been shown to improve growth rate.

Since these mutations likely corresponds to i) adaptation to the SIM medium, ii) are found in clones with very distinct antibacterial activities, iii) appeared in both the E640 and EP640 populations, and iv) are combined with other mutations in the different clones (**Table S3**), we did not test their impact on T6SS function.

### Experimental evolution targeted T6SS promoter and structural components

All the E640 clones with no detectable T6SS antibacterial activity carried mutations within the Sci1 promoter or within genes encoding key structural components of the secretion apparatus, TssM, TssK and TssF. All these mutations were re-introduced in the genome of the ancestral strain, and antibacterial activity assays confirmed that the phenotype was linked to these mutations.

#### Psci1

Two different mutations were identified in the *sci1* promoter (P*sci1*): a 34-bp deletion and a base substitution (**Table 1**). The 34-bp deletion encompasses the -10 transcriptional element (28) and is likely to prevent proper binding of the RNA polymerase. The other mutation, an A➔T substitution, is located 13-bp upstream the *tssB* start codon, and near the *tssB* mRNA ribosome-binding site (RBS) (**Figure 3A**). Structural predictions suggest that the A➔T substitution near the RBS changes the overall 5’UTR mRNA architecture (**Figure 3B**), which may impact mRNA translation. Alternatively, it has been recently shown that a single substitution near the RBS deny the binding and its translation by the ribosome (29,30). In agreement with a potential impact on translation, we observed that the mutation induces a 2-fold reduction of fluorescence in a strain expressing the GFP inserted between the *tssC* and *tssK* genes (transcriptional reporter C-GFP-K, **Figure 3C**).

**Figure 3.**
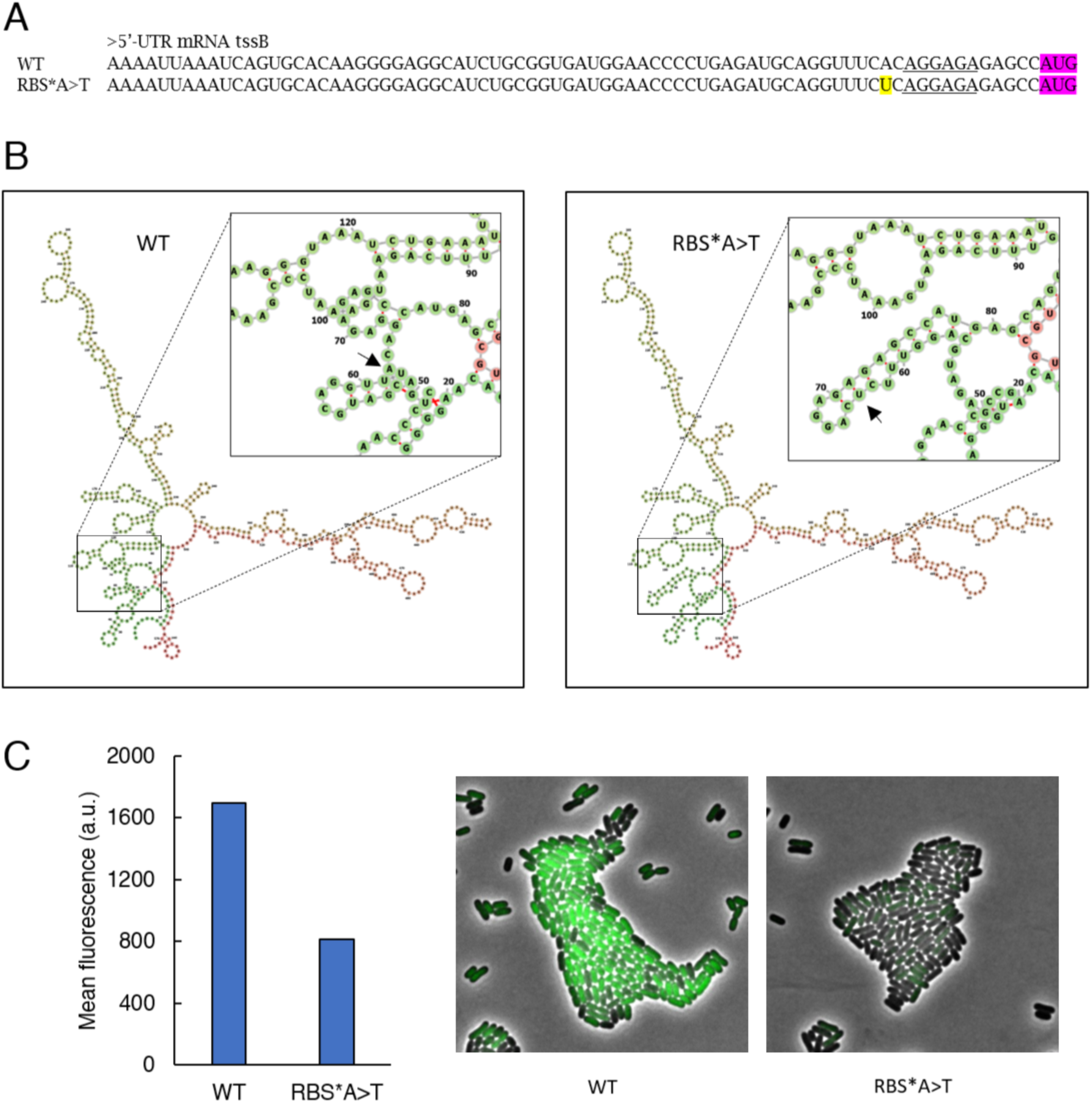
P*sci1* substitution may impact T6SS expression or translation. **A.** Alignment of the *tssB*-mRNA 5’-UTR sequence of the WT and P*sci1*\* evolved clone (RBS*A>T). The consensus RBS sequence (AGGAGA) is underlined. The *tssB* start codon is shown in pink whereas the SNP is highlighted in yellow. **B.** Prediction of the secondary structure of the WT (left) and RBS*A>T (right) *tssB* mRNA using the Vienna RNA Websuite (31). The WT sequence shows interaction between the -13A (arrow) and an upstream nucleotide, whereas the variant -13U (arrow) interacts with the -6A of the RBS, modifying the secondary structure and RBS exposition. **C.** Mean fluorescence of a EAEC 17-2 C-GFP-K population in the WT compared to the RBS*A>T variant (*n*=1100 cells) (left). Representative fluorescence microscopy image of EAEC 17-2 C-GFP-K WT or RBS*A>T (right).

#### TssM

TssM is an inner membrane protein that assembles, together with TssL and TssJ, the T6SS membrane complex in EAEC (32,33). The C-terminal domain of TssM interacts with the TssJ outer membrane lipoprotein in the periplasm (34). The identified mutation is a 1-bp deletion of nucleotide 869, hence producing a frameshift and a premature stop codon removing part of the cytoplasmic domain, the third transmembrane helix and the periplasmic domain (**Figure 4A**). Interestingly, a recent comparative genomic study on multi-drug-resistant *E. coli* lineages identified *tssM* as the target of an insertion sequence in one isolate, likely disrupting TssM function (15). Using bacterial two-hybrid, we showed the loss of interaction between TssM and TssJ when introducing the mutation found in the E640 evolved clone (**Figure 4B**). By avoiding this essential interaction, this mutation prevents T6SS sheath assembly (**Figures 4C**) and antibacterial activity (**Figure 4D**).

**Figure 4.**
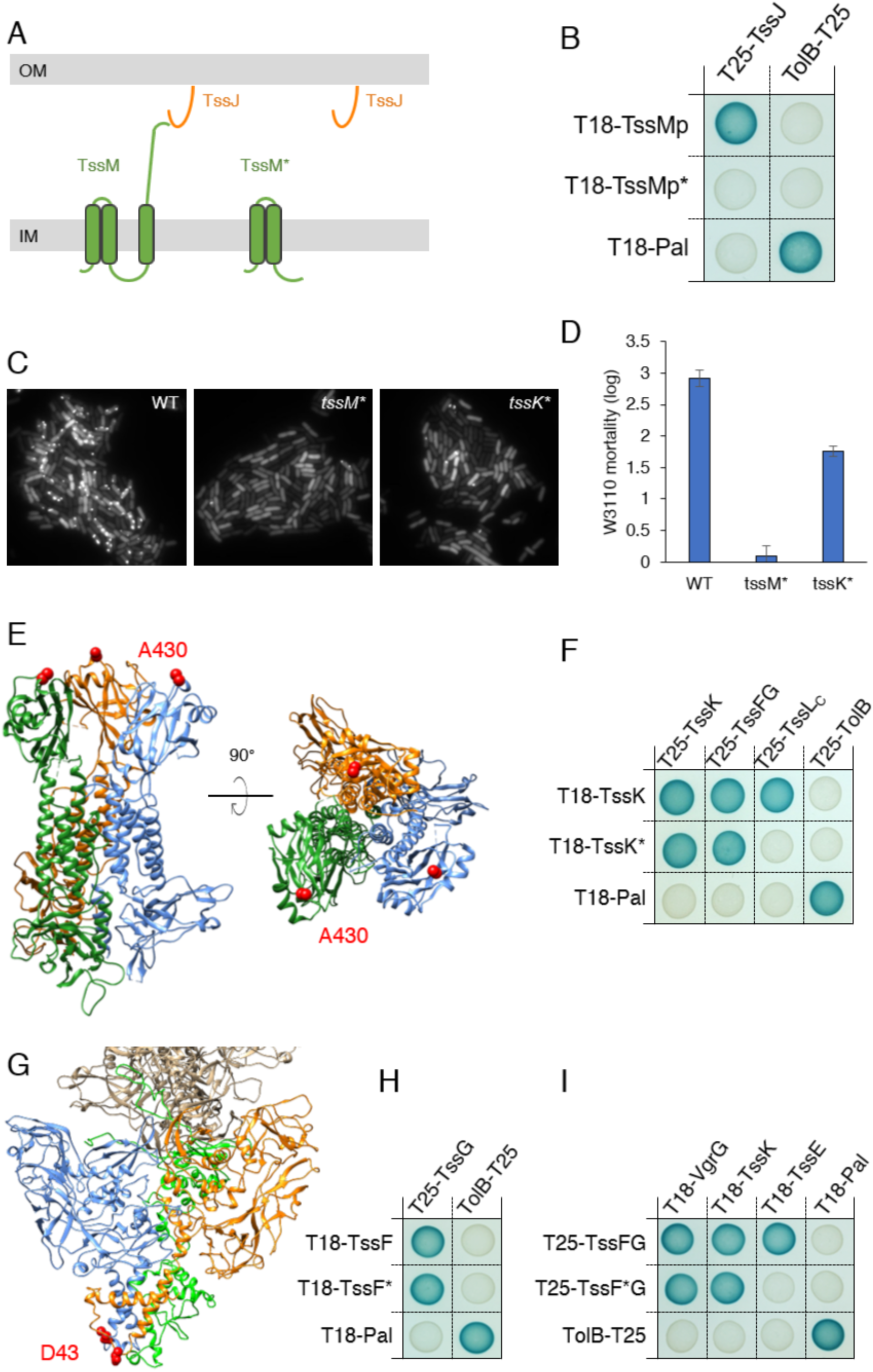
T6SS *tssM*, *tssK* and *tssF* mutations affect the T6SS assembly and activity. **A.** Schematic representation of the TssM protein and its truncated variant (TssM*)(green), and the TssJ OM-associated lipoprotein (orange). IM, inner membrane; OM, outer membrane. **B.** Representative fluorescence microscopy fields of the WT, *tssM\** and *tssK\** strains producing TssB-sfGFP. **C.** Statistical analyses of WT, *tssM\** and *tssK\** cells with polymerized and dynamic sheaths (*n*=1000 cells). **D.** Competition assay between the WT, *tssM\** and *tssK\** strains and the susceptible W3110 strain using the SGK assay (3 independent replicates). **E.** Side (left) and top (right) views of the EAEC TssK trimer structure (pdb: 5M30). Each monomer is shown with a different color, and the position of residue A430 is shown as a red sphere. **F.** Bacterial two hybrid assay. WT TssK or its A430E variant (TssK*) fused to the T18 domain of the adenylate cyclase were tested against TssK, TssFG or the cytoplasmic domain of TssL (TssL_C_) fused to the T25 domain. **G.** Side view of the EAEC TssFGK wedge complex (pdb: 6GIY). TssG is colored green, the two TssK trimers are colored grey whereas the two TssF copies are colored in orange and blue. The position of residue D43 is shown as a red sphere. **H, I.** Bacterial two hybrid assay. WT TssF or its D43H variant (TssF*) were tested against TssG (**H**), and VgrG, TssK and TssE (**I**).

#### TssK

TssK is a trimeric cytoplasmic protein that assembles, together with TssE, -F and -G the T6SS baseplate wedges (35–38). TssK resembles phage-like receptor-binding proteins and anchors the baseplate to the membrane complex via specific interactions between its C-terminal domain and the cytoplasmic domain of the TssL inner membrane protein (35,39–41). The mutation identified in this study corresponds to a substitution of Ala-430, located in one of the loops of the TssK C-terminal domain (**Figure 4E**), downstream residue Pro-429 that establishes intimate contacts with TssL (41). In agreement with these latter study, the TssK A430E substitution did not impact TssK oligomer formation or interaction with TssFG but prevented TssKL complex formation (**Figure 4F**), significantly decreasing T6SS sheath assembly (**Figures 4C**) and antibacterial activity (**Figure 4D**).

#### TssF

TssF forms with TssG the trifurcating unit skeleton of the T6SS baseplate wedge (36–38). The mutation that appeared during experimental evolution corresponds to the substitution of Asp-43 by a histidine residue (**Figure 4G**), which does not impact formation of the TssFG complex (**Figure 4H**) but prevented recruitment of TssE to the TssFG unit (**Figure 4I**).

### Mutations outside the T6SS gene cluster targeted the transcriptional RfaH antiterminator, affecting proper *sci1* transcription

Two mutations located outside the T6SS gene cluster were also identified, affecting the *csgE* and *rfaH* genes. A 1-bp deletion producing a frameshift and a premature stop codon in the *csgE* gene was selected. *csgE* belongs to the *csgDEFG* operon, which, together with the *csgABC* operon, encode the components of surface-associated curli (42). Curli are amyloid fibers that are involved in the adhesion to host epithelial cells by binding fibronectin and MHC-1 (43) and biofilm formation (44,45). CsgE is a periplasmic protein that interacts with the outer membrane protein CsgG to form the secretion canal that exports and assembles the extracellular CsgA and CsgB curli subunit to the cell surface (46–49). Reconstruction of the *csgE* frameshift in the EAEC 17-2 ancestral strain did not reproduce the phenotype observed in the evolved strain, suggesting that *csgE* is not involved in the T6SS activity but rather could be a metabolic adaptation or be required for T6SS-related function in other conditions (**Figure S3**).

The mutation that appeared in the *rfaH* gene is a 4-bp deletion frameshift inducing a premature stop codon. RfaH is a transcriptional regulator that unlocks the RNAP pausing at specific sites named the *ops* elements (consensus GGCGGTAGCGTG) (50). The RNAP-*ops* complex is recognized by RfaH, which binds and stabilizes the RNAP by its N-terminal domain and recruits the ribosome by its C-terminal domain, enhancing the transcription and translation efficiency (51,52). *Ops* elements are usually found in large operons such as those encoding the capsule biosynthesis and secretion machine (*wza*, 21 kb) or secretions systems (T4SS, 33 kb; T1SS, 23 kb) (53). In EAEC, we found the *ops* consensus and derivative sequences in genes related to diverse functions such as transposition, exopolysaccharide production, membrane transporter, LPS biosynthesis or metabolism (**Table S4**). Interestingly, we identified two *ops* elements within the EAEC T6SS *sci1* gene cluster, one with a consensus sequence located in the intergenic region between the *hcp* and *clpV* genes (*ops1*), and a second, less consensus, but with the most conserved GGTAGNNT motif, located in the intergenic region between the accessory genes *EC042_4547* and *EC042_4548* (*ops2*) (**Figure 5A**). Mutagenesis of the *ops1* and *ops2* elements indeed impaired T6SS activity (**Figure 5**). Mutation in *ops1* element completely abolished T6SS assembly (**Figure 5C**) and, consequently, reduced antibacterial activity (**Figure 5D**). This result is consistent with the *ops1* role in governing the proper transcription of the downstream essential structural genes, suggesting a defect in the proper amount of T6SS proteins translated from genes downstream the *ops1* element. In agreement with the position of the *ops2* element, located in the T6SS accessory gene operon, the *ops2* mutation impacted T6SS sheath assembly without abolishing it (**Figure 5C**). To better understand the contribution of *ops* elements and *rfaH* in the transcription of T6SS genes, transcriptional fluorescent reporter strains were engineered, in which the mCherry-coding sequence was inserted at different positions in the T6SS gene cluster in the WT and *rfaH*, *ops1* and *ops2* mutants: upstream the *ops1* element (between the *tssC* and *tssK* genes, C-mCh-K), between the *ops1* and *ops2* elements (between the *clpV* and *EC042_4531* genes, ClpV-mCh-31), and downstream the *ops2* element (between the *EC042_4548* and *EC042_4549* genes, 48-mCh-49). Fluorescence measurements showed that the C-mCh-K reporter remained unchanged in all the strains (**Figure 5E**). In contrast, the transcription of the ClpV-mCh-31 reporter drastically decreased in the *ops1* and *rfaH* strains, and that of the 48-mCh-49 reporter decreased in *ops1*, *ops2*, and *rfaH* strains (**Figure 5E**). Taken together, these results demonstrate that RfaH is required for the proper transcription of the T6SS gene cluster in EAEC, likely by solving transcriptional pause at *ops* sites located within the cluster.

**Figure 5.**
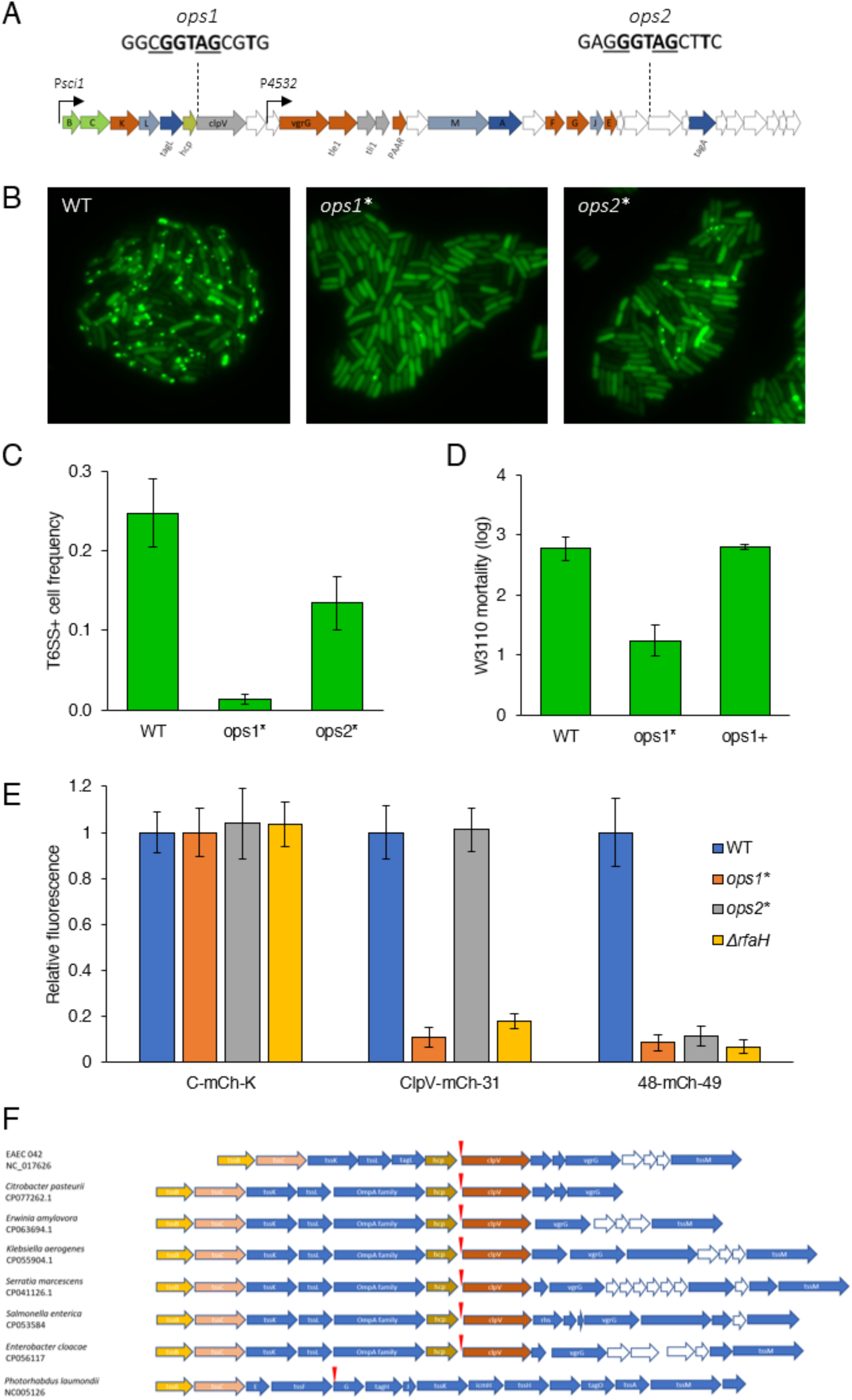
RfaH regulates proper *sci1* expression via conserved *ops* elements. **A.** Schematic representation of the EAEC *sci1* T6SS gene cluster, highlighting the position and the conserved sequence of the two *ops* elements. The most conserved residues are shown in bold whereas the critical residues for RfaH binding are shown underlined. **B.** Representative fluorescence microscopy fields of the WT, *ops1*\* and *ops2*\* strains producing TssB-sfGFP. **C.** Statistical analyses of WT (*n*=2500), *ops1\** (*n*=2500) and *ops2\** (*n*=1300) cells with polymerized and dynamic sheaths. **D.** Competition assay between the WT, *ops1\** and reconstituted *ops1^+^*strains and the susceptible W3110 strain using the SGK assay (3 independent replicates). **E.** mCherry fluorescence measurements in WT (blue bars), *ops1\** (orange bars), *ops2\** (grey bars) and *rfaH* (yellow bars) cells with mCherry insertions between the *tssC* and *tssK* (C-mCh-K), *clpV* and *EC042_4531* (ClpV-mCh-31), or *EC042_4548* and *EC042_4549* (48-mCh-49) genes. The mean fluorescence of each construction was normalized to the WT strain. Standard deviation bars represent the error between 3 biological replicates. **F.** Syntheny of the *ops1* element in T6SS gene clusters in different enterobacteria. The position of a consensus *ops* element is indicated by the red arrow.

As T6SS genes are usually clustered in large operon, we asked whether RfaH-dependent regulation of the T6SS is a conserved mechanism. Genomic analyses on representative enterobacteria showed that *ops1* elements is highly conserved in T6SS clusters (**Figure 5F**). This concerns various Enterobacteriaceae such as human pathogens (*Citrobacter pasteurii* and *Klebsiella aerogenes*), entomopathogens (*Photorhabdus laumondii* and *Serratia marcescens*) and plant pathogens (*Erwinia amylovora*). These *ops* elements possess a high synteny with EAEC *ops1* as they locate in the *hcp*-*clpV* intergenic region in all strains with the exception of *P. laumondii* in which the *ops* element is found between *tssF* and *tssG* genes (**Figure 5F**). The presence of T6SS *ops* sites in diverse clusters suggests that RfaH-dependent transcription of T6SS gene clusters is a conserved mechanism across enterobacteria.

## DISCUSSION

Experimental evolution is a powerful technique to understand the dynamics of emergence of mutations that optimize a process in a specific laboratory-controlled condition (16). It is also a powerful tool to identify new key elements that regulate or modulate a process. Here we conducted an experimental evolution of the EAEC strain 17-2 in competition experiments against susceptible or immune cells in SIM medium, which is the inducing medium for the antibacterial Sci1 T6SS. We observed that while the population evolved in the presence of susceptible cells retains antibacterial activity over 640 generations, about 90% of the cells of the population evolved in the presence of immune cells (EP640) presented a significant attenuation or total loss of T6SS activity.

This result, which demonstrates that cells have a tendency to eliminate the T6SS in conditions in which it is not necessary, i.e., in presence of immune cells or in pure culture, supports the hypothesis that classical laboratory strains of *E. coli*, which are devoid of T6SS gene clusters (or only remnants of them), have evolved over decades of pure cultures in laboratory conditions. Why losing T6SS activity when not necessary? One obvious hypothesis will be that T6SS production, assembly, function or recycling is energetically costly for the cell, as previously shown for the T3SS in *Salmonella enterica* Typhimurium. An experimental evolution of the *S.* Typhimurium *hns* mutant, which is highly impaired in growth under laboratory conditions, led to the systematic deletion of a large portion of the costly SPI-1 T3SS gene cluster which is induced in the *hns* mutant, allowing to counteract the fitness defect of the *hns* deletion (54). However, while the T6SS SPI-6 gene cluster is also induced in the *hns* mutant (55), no mutations impacting the T6SS were identified, suggesting a lower energetic cost for this apparatus. Indeed, recent experiments did not show any significant impact of the T6SS on growth in EAEC (56–59). Thus, why bacteria gain and loss the T6SS over the course of evolution is still elusive.

To identify the genetic modifications that conducted to decreased or loss of T6SS activity, selected clones from the two populations were subjected to whole genome sequencing and the genomes compared to the ancestral strain. Clones with both reduced or no T6SS activity were selected, as we reasoned that they might provide different information. Clones with no activity are likely to bear mutations that directly affect genes that are essential for T6SS assembly or function. In contrast, clones with decreased activity might impact genes encoding accessory subunits of the T6SS or processes that indirectly affect T6SS expression, production, assembly or function (60).

First, we identified mutations common in both conditions suggesting no direct link to competition but rather corresponds to adaptative changes to the medium. Indeed, two mutations targeting the RNA polymerase *rpoC* and ribonuclease *rne* genes, are known adaptative conditions in minimal medium and increase the general fitness in nutrient starvation. Two other mutations, targeting the glycerol transporter *glpK* and the *aqpZ* aquaporin genes, were also found during experimental evolution of *E. coli* or *Salmonella enterica* Typhimurium grown in minimal media with glycerol as carbon source, such as the SIM medium used in our study (17).

The other mutations we specifically identified in the population evolved in the presence of immune cells were found within the *sci1* promoter, within T6SS gene coding sequences (*tssF*, *tssK*, *tssM*) or within genes that are not known to be related to T6SS regulation or function, *rfaH* and *csgE*. To verify that these mutations were responsible for the loss or decrease of T6SS activity, they were introduced into the ancestral strain. With the exception of the *csgE* mutation, which did not show any impact on T6SS activity when reconstructed, all the other mutations phenocopied the activity of their cognate evolved clones.

The two mutations found in the promoter consist to a large deletion encompassing the -10 transcriptional element, likely responsible for an expression defect of the T6SS gene cluster, and to a SNP in the 5’-UTR region, at very close proximity to the RBS. We did not characterize the first deletion since the impact of the -10 transcriptional element deletion on T6SS expression looks obvious regarding the regulation of EAEC Sci1 (61). The second mutation near the RBS is responsible for a decrease in T6SS expression. While we do not have experimental evidence of the molecular mechanism, one may hypothesize that this substitution impacts the mRNA secondary structures that may decrease ribosome access to the RBS, based on predictive tools (29,30). Thus, further analyses are required to understand the impact of this mutation on mRNA structure and stability.

The mutations in the genes encoding the TssF, TssK and TssM subunits were shown to impact the TssF-TssE, TssK-TssL and TssM-TssJ interactions, respectively, which are essential for proper assembly of the T6SS (38). Interestingly, recent phylogenetic analysis reported gain and loss of the T6SS activity during the evolution (15,62–65). *Vibrio cholerae* evolved as pandemic and non-pandemic strains via point mutations in T6SS *clpV* or *tssM* genes (62,64–66). Comparative genomics in *Bacteroidetes* revealed frameshifts caused by independent 2- or 5-bp deletions that affect the *tssC* or *vgrG* genes, suggesting that these strains are in the process of naturally losing the T6SS (67). More generally, laboratory experimental evolutions impose a strong selective pressure on highly conserved protein domain which it is hypothesized to be easily reversed (68). It would be interesting to test whether the mutations we identified in our study could be reversed by subjecting them to a novel experimental evolution in the presence of susceptible recipient cells. More generally, using experimental evolution with diverse strains could be a basis to compare what is observed by comparative genomics and to test whether this approach could reflect the evolution in nature. We have to note that our conditions targeted the antibacterial Sci1 T6SS of EAEC whereas comparative genomic studies can show evolution of other T6SS types potentially targeting other biological forms such as host eukaryotic cells, fungi or protists. Thus, experimental evolution on other T6SS in the appropriate conditions should be perform to support hypothesis on genetic basis for T6SS evolution found in comparative studies.

The most interesting mutation acquired during the experimental evolution in the presence of immune cells is the frameshift within the *rfaH* gene. The RfaH antiterminator is responsible for RNA polymerase stabilization for the transcription of long operons, by resolving RNAP pausing at specific *ops* sequences and suppressing premature termination (51). Interestingly, we found two functional *ops* elements within the Sci1 T6SS gene cluster, located in intergenic regions within the main (*ops1*) and accessory (*ops2*) operons. We further showed that these two elements are important for the transcription of downstream genes, suggesting that they can enhance RNA polymerase transcription processivity. Therefore, RfaH and the two *ops* elements are new regulators of T6SS expression in EAEC. A comparative genomic analysis suggested that the RfaH-dependent regulation of the T6SS is likely to be conserved in enterobacteria, as consensus *ops* sequences are found in intergenic regions in pathogenic and commensal *E. coli*, *Salmonella*, *Citrobacter*, *Enterobacter*, *Klebsiella*, *Serratia* or *Photorhabdus* species. In agreement with this observation, it has recently been showed that the *rfaH* deletion impacted T6SS gene expression in *Erwinia amylovora* (69) and that RfaH solves RNA polymerase pausing and counteracts H-NS silencing of T6SS genes in uropathogenic *E. coli* (70). One may hypothesize that this additional counter-silencer role of RfaH is conserved as T6SS gene clusters in *E. coli* EDL933, *S.* Typhimurium and *Edwardsiella piscicida*, which are regulated by H-NS (71), carry *ops* elements. Further investigations are necessary to understand how RfaH is involved in the regulation of the T6SS, notably by studying its own regulation and going deeper in the biochemical interaction with its DNA, RNA polymerase and ribosome partners. The observation of *ops* elements in several loci on EAEC genome suggests that the T6SS could be co-regulated with other systems and be part of a specific program governed by RfaH. We uncovered a new layer of regulation among transcriptional and post-translational checkpoint in the T6SS field that we need to better characterize to understand how and when T6SS are employed.

Taken together, this study highlights the power of experimental evolution to understand the molecular basis of the evolution of T6SS in bacterial strains but also to unveil new regulatory elements that optimize its expression or function. Further work aiming at characterizing more evolved clones or the archived intermediate evolved populations will likely provide additional information such as new accessory genes or how loss-of-function mutations emerge.

## MATERIAL AND METHODS

### Strains and growth conditions

Bacterial strains and plasmids used in this study are listed in **Table S1**. *Escherichia coli* strains were routinely grown in LB or T6SS inducing medium (SIM: M9 minimal medium supplemented with 0.2% glycerol, 1 μg/ml vitamin B1, 40 μg/ml casaminoacids and 10% (v/v) LB medium) (28) at 37°C. When necessary cultures were supplemented with kanamycin (50 μg/mL), ampicillin (100 μg/mL) or chloramphenicol (40 μg/mL).

### Experimental evolution procedure

The EAEC 17-2 ancestral strain was engineered by replacing the *lacZ* gene by the kanamycin resistance (Kan^R^) cassette from pKD4 by λ-red-mediated recombination (72) using plasmid pKOBEG (73)to yield strain EAEC 17-2 Δ*lacZ*Ω*kan*. Serial passage experiments were conducted in two conditions, in the presence of the W3110 p*mCherry*-ampR susceptible recipient (EP) or of the EAEC 17-2 immune recipient (E). Strains were cultured in SIM at 37°C with agitation until OD 1 prior to be mixed in 1:4 (v/v) donor:recipient ratio, diluted 1/1000 in 10 mL of fresh SIM, cultured at 37°C with shaking until absorbance at λ=600 nm (*A*_600_) of 0.25, sedimented and further incubated without shaking until an *A*_600_ of 1. Then 200 µL of the co-culture were isolated on LB agar supplemented with kanamycin to select the evolved donor and X-gal to check its phenotype. After ON incubation at 37°C, evolved donor cells were recovered in SIM, the *A*_600_ was adjusted to 1, and cells were archived by storage at -80°C in glycerol and used to seed a new co-coculture by 1/1000 dilution with new susceptible (EP) or immune € cells (Fig. 1).

### Bacterial competition assay

The LAGA and SGK methods were employed to test T6SS activity (74). The LAGA assay is based on the degradation of Chloro-Phenol-Red-β-D-galactopyrannoside (CPRG, yellow color), a membrane-impermeant substrate of the β-galactosidase, which is degraded in Chloro-Phenol-Red (CPR, purple color) by the β-galactosidase released during recipient lysis. The color is therefore a qualitative indication of recipient cell lysis. The SGK assay is based on the lag time necessary for surviving recipient cells to reach a certain *A*_600_, which is inversely proportional to the level of surviving cells, and hence proportional to the antibacterial activity of the donor. For both, donor and recipient strains were precultured in LB overnight and diluted 1/100 in SIM until *A*_600_ reaches 0.8. Donor and recipient were mixed in a 4:1 ratio (v/v) and 10 µL of the mix was spotted on SIM agar plate in triplicate. Spots were dried and incubated at 37°C for 4 hours. For the LAGA qualitative assay, 10 µL of CPRG 2 mM were added on each competition spot. For the SGK assay, each spot were resuspended in 1 mL of selective medium for the recipient and 100 µL of 1/10 dilutions were added to 100 µL of selective medium in a 96-well microplate (Thermofisher, Nunc). Cultures were grown for 15 hours at 37°C with agitation and *A*_600_ was measured every 5 minutes using a microplate reader (TECAN). Serial dilutions of the recipient alone were cultured to establish a standard curve. The time of emergence (Te, time necessary to reach an *A*_600_ of 0.4) of each standard samples was associated with a log difference and a linear regression with equation Δ*log* = *a* × *Te* + *b* was generated. The log difference between each strain was then calculated with their Te using the linear regression equation. All values of each replicate of three biological replicates were compared between strains with a Wilcoxon statistical test using the R software. For both assays, controls with the recipient alone, the donor alone, or the recipient with the ancestral EAEC 17-2 Δ*lacZ*Ω*kan* strain, were systematically performed.

### Whole genome sequencing and bioinformatic analysis

Eighteen evolved clones were selected based on their antibacterial activity and the absence of major growth defects for whole genome sequencing. Genomic DNA was extracted with the DNA MiniKit (Qiagen) and subjected to Illumina sequencing (MicrobesNG, Birmingham, UK). The sequencing files were first merged in fasta format using the Galaxy online plateform (75) and implemented in MAUVE (76,77) to identify SNPs and gaps. mRNA secondary structure predictions were performed by inferring the 5’-UTR-*tssB* mRNA sequence to the RNAfold webserver from Vienna RNA Websuite (31).

### Molecular biology

Oligonucleotides used in this study are listed in **Table S2**. SNPs and gaps were reintroduced in the ancestral EAEC 17-2 wild-type strain by chromosomal directed mutagenesis using the thermosensitive pKO3 suicide vector (78). Briefly, a 1-kb DNA fragment comprising the site to be mutagenized was amplified by PCR and cloned into the pJET1.2/blunt vector. The region was then mutagenized by Quick-change mutagenesis using the Q5 Pfu polymerase and complementary oligonucleotides bearing the desired mutation or deletion. After verification by DNA sequencing (Eurofins genomics), the mutated fragment was amplified by PCR and cloned into the pKO3 plasmid. After verification by DNA sequencing, the pKO3 recombinant suicide vector was introduced into EAEC 17-2 by electroporation. The first event of recombination was selected on chloramphenicol LB agar plates at 37°C, and excision of the plasmid by a second recombination event was selected on LB agar plates supplemented with Sucrose 10%. The mutation was checked by DNA sequencing after PCR amplification of the region of interest (Eurofins Genomic). Fluorescent reporter (mCherry or sfGFP) cassettes were amplified by PCR using vectors pKD4-Nter-sfGFPr and pKD4-Nter-mCherry (39) and oligonucleotides carrying 5’ 50-bp extensions corresponding to sequences upstream and downstream the site of insertion, and inserted on the chromosome by λ-red-mediated recombination using plasmid pKOBEG.

### Microscopy analysis

EAEC 17-2 producing the fluorescent reporters were precultured in 3 mL LB overnight, diluted in SIM to an *A*_600_ of 0.8. Cells were concentrated to an *A*_600_ of 8 by centrifugation and resuspension, and 1 µL of the suspension was spotted on SIM-agarose pads casted using a gene frame stuck on a microscopy slide. A high precision cover slip was added prior to the observation using a Nikon Ti2 microscope. A mask was generated from microscopy images using the MiSiC program (79), implemented into the Fiji software (80), and cells were segmented using MicrobeJ plugin (81). Fluorescence intensity for each cell were extracted and the presence of a sheath was determined for a ratio of max fluorescence/mean fluorescence >2 since sheaths are brighter objects in a cell.

## ACKNOWLEDGEMENTS

We thank members of the Cascales laboratory for discussions, and Moly Ba, Isabelle Bringer, and Annick Brun for technical assistance. This work was supported by the CNRS, the Aix-Marseille Université and by grants from the Agence Nationale de la Recherche (ANR-20-CE11-0011), the Fondation pour la Recherche Médicale (DEQ20180339165) and the Fondation Bettencourt-Schueller to EC.

## Data availability

All data are shown in the manuscript or in the supplemental material.

