## Supplemental Information for "Molecular bases for the loss of type VI secretion system activity during enteroaggregative *E. coli* experimental evolution"

**Table S1. Strains used in this study**

| Name | Strain | Description | Source |
|---|---|---|---|
| **EAEC 17-2** | Entero-aggregative *E. coli* 17-2 | Wild-type strain, predator | |
| *sci1* | 17-2 Δ*sci1* | T6SS-1 deficient | |
| *lacZ* | 17-2 *lacZ*::kan | Ancestral strain carrying KanR for EE | This study |
| *sci1 lacZ* | 17-2 Δ*sci1 lacZ*::kan | | This study |
| *tagA* | 17-2 Δ*tagA* | Intermediate phenotype + | Santin et al., 2018 |
| *clpV* | 17-2 Δ*clpV* | Intermediate phenotype - | Douzi et al., 2016 |
| EP-640 | 17-2 *lacZ*::kan evolved with preys | EP-640 evolved population | This study |
| E-640 | 17-2 *lacZ*::kan evolved without preys | E-640 evolved population | This study |
| B-GFP | 17-2 TssB-GFP | T6SS-1 translational reporter | |
| C-GFP-K | 17-2 *tssC-gfp-tssK* | T6SS-1 transcriptional reporter | This study |
| *ops1* | 17-2 *ops1*-mut | | This study |
| *ops1*1* | 17-2 *ops1*-mut reversion | | This study |
| *ops*2 | 17-2 *ops2*-mut | | This study |
| EE-F1 | 17-2 Δ*rfaH* | | This study |
| *csgE* | 17-2 Δ*csgE* | | This study |
| *tssM* | 17-2 Δ*tssM* | | This study |
| *tssK* | 17-2 $TssK_{A430D}$ | | This study |
| RBS* | 17-2 P*sci1*-EE | | This study |
| W3110 | *E. coli* K-12 W3110 | Prey strain | |
| W3110 mCherry | W3110 p-mCherry+AmpR | Prey carrying plasmid | |
| pJET | AmpR | Cloning vector for quick change | |
| pKO3 | SacB, CmR, 30°C thermosensitive | Cloning vector for directed mutagenesis | |
| pKD4 | FRT-KanR-FRT | Template for lambda-red recombination | |
| pKOBEG | Rec genes, CmR, 30°C thermosensitive | Plasmid-encoding lambda-red system | |
| pBAD33 | Plac, CmR | Expression vector for complementation | |

**Table S2. Oligonucleotides used in this study**

| Cloning pJET/pKO3 | Sequence |
|---|---|
| 5-pKO3-ops1 | TATCGGATCCggctgaaagatgatggcg |
| 3-pKO3-ops1 | ACGGTCGACcggtatacctgccgtatcc |
| 5-pKO3-ops2 | TATCGGATCCcgtttggtgaggatgtc |
| 3-pKO3-ops2 | ACGGTCGACcctgctgaatatccataattttg |
| 5-pKO3-rfaH | TATCCCCGGGcatcgccagagacg |
| 3-pKO3-rfaH | ACGGTCGACgccatcatcccggc |
| 5-pKO3-csgE | TATCGGATCCcggataaaaagcttatcc |
| 3-pKO3-csgE | ACGGTCGACgggtgtttcaatacc |
| 5-pKO3-glpK | TATCCCCGGGcggacgatggcaacgg |
| 3-pKO3-glpK | ACGGTCGACggcgatcccggagattgg |
| 5-pKO3-aqpZ | TATCGGATCCggtggatatgttcag |
| 3-pKO3-aqpZ | ACGGTCGACcccagtccttgcaaag |
| 5-pKO3-rpoC | TATCGGATCCcgcacaccatgccgg |
| 3-pKO3-rpoC | ACGGTCGACgcgtcttctgcagtcacc |
| 5-pKO3-tssM | TATCGGATCCgggacgtaagatacgca |
| 3-pKO3-tssM | ACGGTCGACcctgagctgctgctc |
| 5-pKO3-tssF | TATCGGATCCggtgttgtaataagctgatttg |
| 3-pKO3-tssF | ACGGTCGACgggtgaaaaccatccgtc |
| 5-pKO3-tssK | TATCGGATCCccctgaacagtgcggag |
| 3-pKO3-tssK | ACGGTCGACcggtgcacgcaccac |
| 5-pKO3-rne | TATCGGATCCggcgacgcgactagac |
| 3-pKO3-rne | ACGGTCGACcctgacgcaccgcttc |
| 5-pKO3-Psci | TATCGGATCCGCCGGCAGGGATTTCG |
| 3-pKO3-Psci | ACGGTCGACCGGCATGCGCTTCAG |
| **Quick-change** | |
| 5-pKO3-OPS1-mut | cctgggttacggcgaagggcaaaagcgtgcctgacggtctcctg |
| 3-pKO3-OPS1-mut | caggagaccgtcaggcacgcttttgcccttcgccgtaacccagg |
| 5-pKO3-OPS2-mut | atgacgcccggggagaggCgTGTTgcGAGTcgactggagagct |
| 3-pKO3-OPS2-mut | agctctccagtcgACTCgcAACAcGcctctccccgggcgtcat |
| 5-pKO3-rfaH-del | atgcaatcctggtatttacttgcaagcgcgggcaacttca |
| 3-pKO3-rfaH-del | tgaagttgcccgcgcttgcaagtaaataccaggattgcat |
| 5-pKO3-csgE-del | tctattggccatgatttttacgagcctttagtgataaatg |
| 3-pKO3-csgE-del | catttatcactaaaggctcgtaaaaatcatggccaataga |
| 5-pKO3-glpK-A206T_D69V | agaagtgctggcgaaagccgttatcagttccgatcaaattg |
| 3-pKO3-glpK-A206T_D69V | caatttgatcggaactgataacggctttcgccagcacttct |
| 5-pKO3-aqpZ-G564A_A188A | aacacttctgttaacccggcacgcagcaccgcggttgctat |
| 3-pKO3-aqpZ-G564A_A188A | atagcaaccgcggtgctgcgtgccgggttaacagaagtgtt |
| 5-pKO3-rpoC-del | ctcaacgtgttcgaaggtgaacgtggtgacgtaatttccg |
| 3-pKO3-rpoC-del | cggaaattacgtcaccacgttcaccttcgaacacgttgag |
| 5-pKO3-tssM-del | agagcgattcagacgaaatatgatgctgtggtggagccat |
| 3-pKO3-tssM-del | atggctccaccacagcatcatatttcgtctgaatcgctct |
| 5-pKO3-tssF-G127C_D43H | tcgataaagcgggtacgcctcatccctgcgtggaacgcctg |
| 3-pKO3-tssF-G127C_D43H | caggcgttccacgcagggatgaggcgtacccgctttatcga |
| 5-pKO3-tssK-C1289A_A430E | ttgtactttctacaccccggaatcgctgggagatgtgaaac |

| | |
|---|---|
| 3-pKO3-tssK-C1289A_A430E | gtttcacatctcccagcgattccggggtgtagaaagtacaa |
| 5-pKO3-rne1-G514A_G172S | ggcttatcgtgcgcaccgctagcgtcggcaaatctgctgag |
| 3-pKO3-rne1-G514A_G172S | ctcagcagatttgccgacgctagcggtgcgcacgataagcc |
| 5-pKO3-rne2-G515C_G172A | gcttatcgtgcgcaccgctgccgtcggcaaatctgctgagg |
| 3-pKO3-rne2-G515C_G172A | cctcagcagatttgccgacggcagcggtgcgcacgataagc |
| 5-pKO3-Psci1-A-T_EE | acccctgagatgcaggtttctcaggagagagccatgagcag |
| 3-pKO3-Psci1-A-T_EE | ctgctcatggctctctcctgagaaacctgcatctcaggggt |
| 5-pKO3-Psci1-del_EE | atttttcagatcttcgtcctgggaggcatctgcggtgatg |
| 3-pKO3-Psci1-del_EE | catcaccgcagatgcctcccaggacgaagatctgaaaaat |
| 5-pKO3-Psci(-26A-T) | ctgcggtgatggaacccctgTgatgcaggtttcacaggaga |
| 3-pKO3-Psci(-26A-T) | tctcctgtgaaacctgcatcAcaggggttccatcaccgcag |
| 5-pKO3-Psci(-26A-T)(-13A-T) | ctgcggtgatggaacccctgTgatgcaggtttcTcaggaga |
| 3-pKO3-Psci(-26A-T)(-13A-T) | tctcctgAgaaacctgcatcAcaggggttccatcaccgcag |
| **Cloning pBAD33** | |
| 5-pBAD33-csgE | attgagctcaggaggtattacaccAtgaaacgttatttacgctggattg |
| 3-pBAD33-csgE | agtgtcgacttagaattcatcatgcgccaaatc |
| 5-pBAD33-rfaH | attgagctcaggaggtattacaccatgcaatcctggtatttac |
| 3-pBAD33-rafH | agtgtcgacttagagtttgcggaactcgg |

### Table S3. Lineages during the EE

| Clones | Mutated genes | | | | | | | T6SS activity |
|---|---|---|---|---|---|---|---|---|
| B6-C2 | *rpoC* | | | | *Psci-SNP* | | | + |
| A111-E7-F9 | *rpoC* | | | | *tssF* | | | + |
| G7-E1-C1-E5 | *rpoC* | *rne2* | | | *tssM* | | | - |
| H7-D8 | *rpoC* | *rne2* | | | | *csgE* | | = |
| C12 | | | *glpK* | *aqpZ* | | | | + |
| F1 | | | *glpK* | *aqpZ* | | | *rfaH* | - |
| G5-A112 | | | *glpK* | *aqpZ* | *tssK* | | | - |
| D1-E8-H8 | | *rne1* | *glpK* | *aqpZ* | *Psci-Del* | | | - |

Left column indicates the name of the clone. Middle columns indicate the metabolic (blue), functional (yellow) or other potentially functional (red) genes mutated. Positive, neutral, or negative T6SS activity are referred to (+), (=) or (-), respectively.

**Table S4. Distribution of *ops* consensus and arbitrary variations in the EAEC 042 genome**

| ***ops*** **sequence** | **Gene in or downstream** |
|---|---|
| **GGCGGTAGCGTG** | |
| **570 711** | *ushA* |
| **2 375 242** | EC042 RS12205 (glycosyltransferase) |
| **2 399 558** | EC042 12310 (EPS) |
| **3 406 147** | EC042 28315 (transposase) |
| **4 188 594** | *rfaQ* |
| **4 859 296** | *hcp*/*clpV* intergenic region (*sci1*) |
| **GGCGGTAGCGTA** | |
| **264 452** | *hcp*/*clpV* intergenic region |
| **671 457** | EC042 RS03115 |
| **1 652 782** | EC042 RS08380 (PAAR) |
| **3 458 321** | EC042 RS17310 (permease) |
| **4 193 660** | *coaBC* |
| **4 647 706** | *birA* |
| **GGCGGTAGCGTT** | |
| **1 017 143** | EC042 RS05010/*artP* |
| **2 780 263** | *ligA* |
| **GGCGGTAGCGTC** | |
| **2 046 382** | EC042 RS10370 |

The sequence variant is indicated by letters in the left columns. Associated numbers in the column correspond to the genome position.

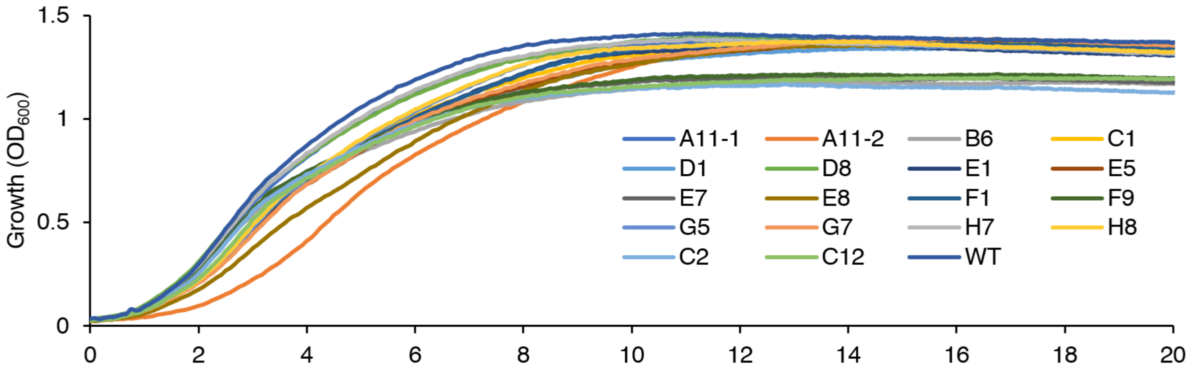


**Figure S1.** Growth curves of evolved clones and the ancestral strain in SIM at 37°C. The graphs represent the average $OD_{600}$ of 15 replicates from 5 independent cultures.

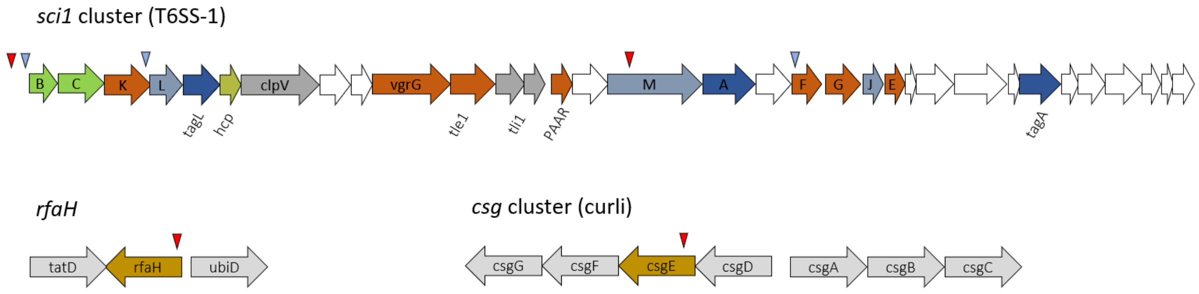


**Figure S2. Schematic representation of functional mutations detected on the genomes.** The arrow indicates a gene and its orientation. The triangle indicates the position of the mutation (red=deletion; blue=SNP).

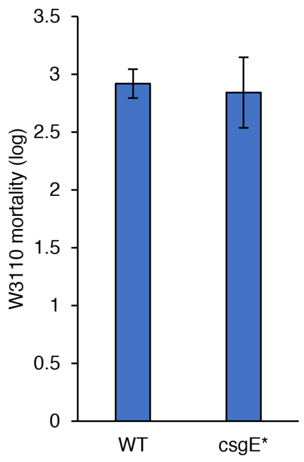


**Figure S3.** Competition assay between EAEC or *csgE** attackers and W3110 recipient cells. Recipient mortality was measured by the SGK method (results from 3 independent replicates).
